# Modular cell-free expression plasmids to accelerate biological design in cells

**DOI:** 10.1101/2020.07.22.216267

**Authors:** Ashty S. Karim, Fungmin Eric Liew, Shivani Garg, Bastian Vögeli, Blake J. Rasor, Aislinn Gonnot, Marilene Pavan, Alex Juminaga, Séan D. Simpson, Michael Köpke, Michael C. Jewett

**Affiliations:** Department of Chemical and Biological Engineering, Northwestern University, Evanston, IL 60208, USA; Chemistry of Life Processes Institute, Northwestern University, Evanston, IL 60208, USA; Center for Synthetic Biology, Northwestern University, Evanston, IL 60208, USA; LanzaTech Inc., 8045 Lamon Ave Suite 400, Skokie, IL 60077, USA; Simpson Querrey Institute, Northwestern University, Chicago, IL 60611, USA

**Author notes:** These authors contributed equally to this work (Ashty S. Karim, Fungmin Eric Liew). Corresponding Authors: Michael C. Jewett, and Michael Köpke.

## Abstract

Industrial biotechnology aims to produce high-value products from renewable resources. This can be challenging because model microorganisms—organisms that are easy to use like *Escherichia coli*—often lack the machinery required to utilize desired feedstocks like lignocellulosic biomass or syngas. Non-model organisms, such as *Clostridium*, are industrially proven and have the desired metabolic features but have several hurdles to mainstream use. Namely, these species grow more slowly than conventional laboratory microbes and genetic tools for engineering them are far less prevalent. To address these hurdles for accelerating cellular design, cell-free synthetic biology has emerged as an approach for characterizing non-model organisms and rapidly testing metabolic pathways *in vitro*. Unfortunately, cell-free systems can require specialized DNA architectures with minimal regulation that are not compatible with cellular expression. In this work, we develop a modular vector system that allows for T7 expression of desired enzymes for cell-free expression and direct Golden Gate assembly into *Clostridium* expression vectors. Utilizing the Joint Genome Institute’s DNA Synthesis Community Science Program, we designed and synthesized these plasmids and genes required for our projects allowing us to shuttle DNA easily between our *in vitro* and *in vivo* experiments. We next validated that these vectors were sufficient for cell-free expression of enzymes, performing on par with the previous state-of-the-art. Lastly, we demonstrated automated six-part DNA assemblies for *C. autoethanogenum* expression with efficiencies ranging from 68-90%. We anticipate this system of plasmids will enable a framework for facile testing of biosynthetic pathways *in vitro* and *in vivo* by shortening development cycles.

## Introduction

Industrial biotechnology often seeks to produce chemical products from inexpensive and prevalent feedstocks, such as lignocellulosic biomass and syngas.^1-3^ While most synthetic biologists work with model organisms like *Escherichia coli* and *Saccharomyces cerevisiae* due to their ease-of-use, these organisms can be limited by accessible feedstocks, products, and stable operating environments in which to work. For example, these organisms do not naturally possess the metabolic pathways required to access the carbon in syngas; rather, researchers turn to diverse genera of non-model organisms and pathways capable of these unique biochemical transformations.^4^ One such genus is *Clostridium*, which includes the cellulolytic *C. thermocellum* as well as the gas-fermenting, acetogenic *C. autoethanogenum*.^5-7^ Despite their utility for biotechnology and commercial deployment, these species grow more slowly than conventional laboratory microbes, are obligate anaerobes, and genetic tools for engineering them are still developing and far less prevalent.

Developments in cell-free synthetic biology have sought to characterize non-model organisms^8-10^ and rapidly test metabolic pathways *in vitro*.^11-14^ By using cell-free gene expression (CFE)^15^ to produce enzymes directly *in vitro*, metabolic pathways can be tested without the need to re-engineer organisms or construct new DNA elements between each engineering cycle.^13,16- 20^ This approach benefits from the ability to test more enzyme variants, the ability to precisely tune reaction conditions and enzyme concentrations, and shorter engineering cycles to down-select promising candidate pathways for *in vivo* biochemical production.^21^ While cell-free pathway prototyping is carried out in a mix-and-match fashion,^13^ cellular expression requires assembly into operons. Additionally the specialized plasmids for *Clostridium* expression and those for CFE are not inherently compatible, for example require different promoters (cell-free expression typically relies the orthogonal T7 system) and additional elements such as Gram-positive replication origin, specific antibiotics cassettes and a low GC content.^22^ This means *Clostridium* optimized DNA for successful pathway designs identified *in vitro* must be separately synthesized and cloned prior to transformation in *Clostridium*, adding several weeks of effort and considerable costs. Streamlining this process would increase the ability to engineer non-model organisms for metabolic engineering applications.

In this work, we present a modular plasmid system on the basis of standard cell-free vector pJL1^*10*,19,23,24^ and universal *Clostridium* shuttle vector system pMTL80000^22^ to rapidly bridge cell-free prototyping efforts and strain engineering in *C. autoethanogenum* and reduce the overall engineering cycle time. Engineering compatible expression plasmids requires fine tuning to minimize impacts on the genetic context of open reading frames, particularly around the critically important ribosome binding site.^13, 19^ First, we designed several plasmid architectures that resemble our top-performing cell-free expression vector, pJL1 (Addgene #69496), and would enable flexible arrangement of genes and promoters (Golden Gate assembly^25^ compatibility) for expression in *C. autoethanogenum*. Next, we validated that these new vectors are sufficient for CFE of biosynthetic enzymes. Then, we demonstrated DNA assembly efficiency ranging from 68-90% when assembling up to six parts for *C. autoethanogenum* expression. We finally showed automation of the whole workflow on two different automation systems. This modular ‘Cell-free to *Clostridium*’ vector system along with high-throughput and automatable workflows will accelerate strain development efforts for *C. autoethanogenum* or other *Clostridium* species and possibly other non-model organisms, by decreasing delays in the transition between cell-free prototyping and cellular validation.

## Materials and Methods

### Strains and Plasmids

For generation of the ‘Cell-free to *Clostridium*’ vector system and cloning, *E. coli* strain TOP10 (Invitrogen) was used. First, the counter-selectable marker *ccdB* (flanked with BsaI recognition sites) was cloned into pMTL82251 and pMTL83151^22^ to generate the recipient *Clostridium* expression vectors (pCExpress). The construction of vectors pD2 and pD4 involved TOPO (Invitrogen) cloning of terminator and promoter parts (flanked with BsaI recognition sites) amplified or synthesized by JGI into the plasmid, pCR-blunt (Invitrogen). The ‘Cell-free to *Clostridium*’ vectors were derived from pJL1 plasmid (Addgene #69496), modified in the T7 promoter region to contain a BsaI recognition site between the RBS and START codon in three variations to generate pD1, pD3, and pD5. All recipient and donor vectors were verified by DNA sequencing.

DNA codon-optimized genes for *C. autoethanogenum* were generated using LanzaTech’s in-house codon optimization software. *E. coli* adapted sequences were generated using codon optimization tools from Twist Biosciences (California, USA). Genes of interest were provided by JGI in the ‘Cell-free to *Clostridium*’ vectors pD1, pD3, and pD5. All vector DNA sequences used in this study are listed in **Supplementary Table 1**, and all DNA parts are listed in **Supplementary Table 2**. The 58 modular vectors containing parts from **Supplementary Table 2** are listed in **Supplementary Table 3**. The biosynthetic genes used in cell-free assays are listed in **Supplementary Table 4**, and those used in GG assembly are listed in **Supplementary Table 5**.

### Cell-free assays

All cell extracts for CFE were prepared with *Escherichia coli* BL21 Star(DE3) (NEB).^21^ These cells were grown, harvested, lysed, and prepared using previously described methods.^19,26^ CFE reactions were performed to express each enzyme individually using a modified PANOx-SP system described in previous pubications.^27,28^ Protein measurements were taken after 20 h. Active super-folder GFP (sfGFP) protein yields were quantified by measuring fluorescence. Two microliters of CFE reaction was added in the middle of the flat bottom of 96-well half area black plates (Costar 3694; Corning Incorporated, Corning, NY). sfGFP was excited at 485 nm while measuring emission at 528 nm with a 510 nm cutoff filter. The fluorescence of sfGFP was converted to concentration (μg/mL) according to a standard curve.^29^ All other proteins were measured using CFE reactions with radioactive ^14^C-Leucine (10 µM) supplemented for incorporation during protein production. We used trichloroacetic acid (TCA) to precipitate radioactive protein samples. Radioactive counts from TCA-precipitated samples was measured by liquid scintillation to then quantify soluble and total yields of each protein produced as previously reported (MicroBeta2; PerkinElmer).^27,30^

### Golden Gate assembly using manual workflow

Two- to six-part DNA assemblies were performed using GeneArt Type IIs (BsaI) assembly kit (Invitrogen, CA). Specifically, 75 ng of recipient vector was used. Other parts (pD1, pD2, pD3, pD4, and pD5) were added in 1:1 molar ratio with respect to the recipient vector along with the GeneArt Type IIs enzyme mix. The reaction was then incubated in a thermocycler (37 °C for 1 min, 16 °C for 1 min, cycled 30X, followed by cooling at 4 °C). Afterwards, the assembly mixture was transformed into *E. coli* Top 10 chemically competent cells (ThermoFisher Scientific, CA), and plated onto LB agar containing appropriate antibiotics. Resulting colonies were screened via PCR for presence of the parts cloned, followed by sequence confirmation via NGS.

### Golden Gate assembly using automated workflow

Two automated assembly workflows were developed, either using the liquid handling robot Hamilton STARLet or the Labcyte Echo 525.^31^ The assembly reactions were carried out with the final concentration for each individual DNA part was 2 nM.^25^ The assembly reaction volume for Hamilton STARLet was 20 µL prepared as follows: 2 µL of each DNA part (10 nM), 10 µL GeneArt Type IIs Assembly Kit BsaI (Invitrogen A15917), and deionized water to a total of 20 µL. When using the Labcyte Echo 525, the reaction volume was downsized to a final volume of 2 µL. All DNA samples were quantified by absorbance at 280 nm, employing a NanoDrop 2000 spectrophotometer (Thermo Fisher Scientific). Reactions were incubated in an INHECO heat block using the following parameters: 37 °C for 2 h, 50 °C for 5 min, 80 °C for 10 min, then stored at −20 °C until transformation. Transformations were also performed using the INHECO blocks: 2 µL of each reaction mix was added to 20 µL of Invitrogen One Shot Top10 chemically competent cells (C404003) and incubated for 20 min at 4 °C. Cells were then heat-shocked at 60 °C for 45 sec, then recovered for 2 min at 4 °C. Afterwards, 180 µL of Super Optimal Broth (Invitrogen 15544034) with catabolite repression (SOC) media was added to cell mixtures, and cells were recovered at 37 °C for 2 h. Finally, 7 µL of each transformation reaction consisting of undiluted culture volume was plated on Lennox Lysogeny Broth (LB) + agar plates containing the appropriate antibiotic and incubated overnight at 37 °C. We randomly chose two colonies to sequence throughout the assembled regions. NGS sequencing confirmed that more than 90% of the clones screened showed complete assemblies.

## Results and Discussion

### Design of a modular ‘cell-free to *Clostridium*’ vector system

Our goal was to develop a DNA vector system that would enable easy exchange of DNA between cell-free plasmids and cellular plasmids. This cell-free to *Clostridium* framework would minimize repetitive DNA synthesis and subcloning, allowing for facile and rapid testing of biosynthetic pathways *in vitro* and *in vivo* (**Figure 1A**). To achieve this goal, CFE expression plasmids were modified via addition of Golden Gate (GG) sites so that these can be used directly for multiple DNA parts assembly directly into *Clostridium* expression vectors (**Figure 1B**). With such a system in place, genes could be ordered once with *Clostridium* codon-adapted sequences in these plasmids, prototyped in cell-free reactions using these plasmids, and then the best performing gene variants could be assembled from the same plasmids into *in vivo* expression plasmids via one-step GG assembly.

**Figure 1.**
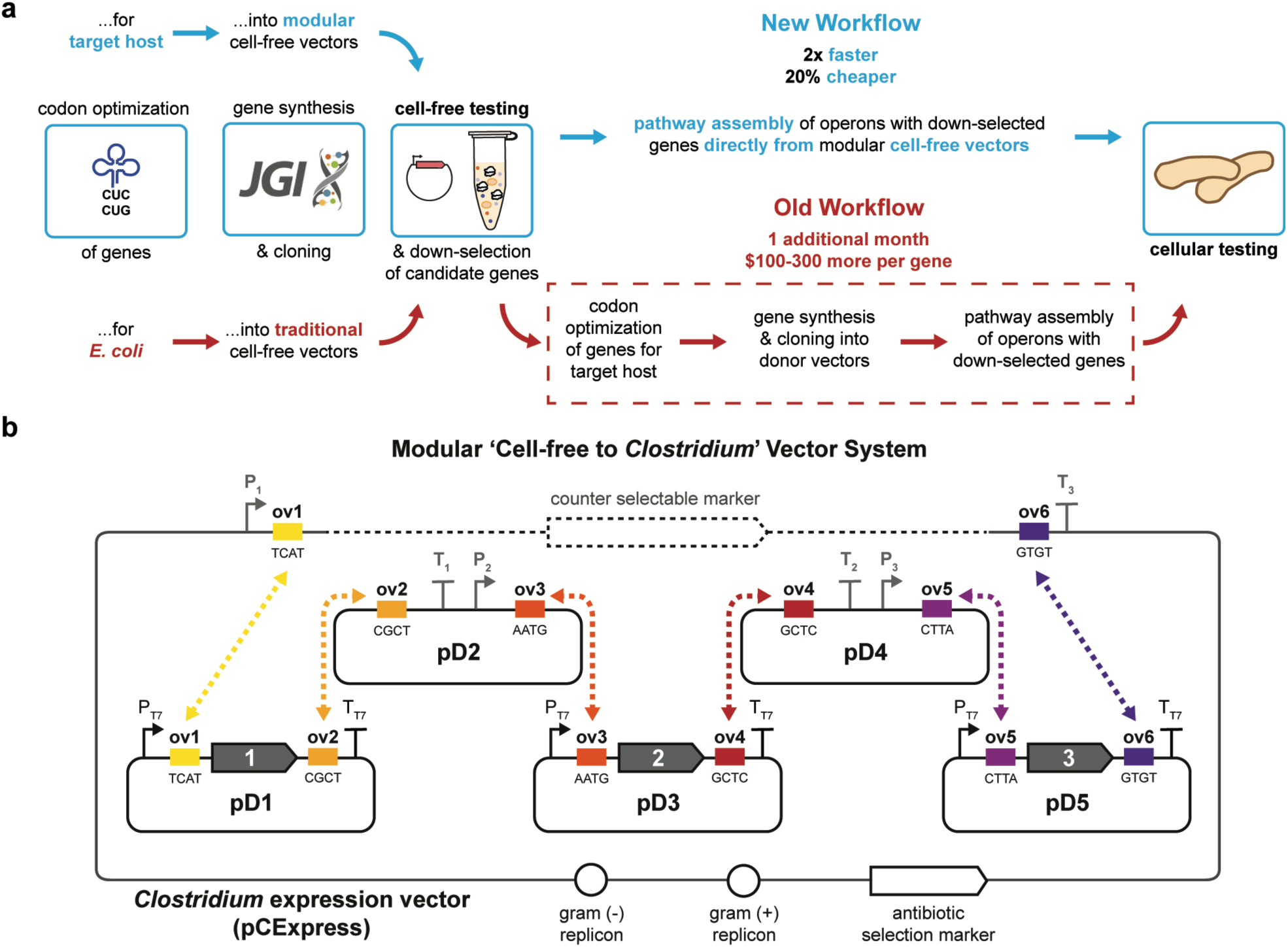
A framework for a modular ‘cell-free to *Clostridium*’ vector system that enables seamless assembly of cell-free vectors into a *Clostridium* expression vector. (a) A schematic representation of how information between *in vitro* and *in vivo* needs is used to design DNA sequences, JGI facilities can construct DNA designs, and DNA materials can be used in both *in vitro* and *in vivo* experiments. (b) The architecture of the modular vector system is shown. Cell-free vectors are made compatible for assemblies by adding unique overhang (Ov) sites generated from *Bsa*I digests.

As a starting point, we designed six total vectors. Three vectors, pD1, pD3, and pD5, were constructed by adding GG sites (BsaI recognition sites) within the T7 promoter region of the pJL1 vector (Addgene #69496), a standard CFE expression plasmid. These vectors were designed to serve as gene donor vectors for assembly in a new recipient vector based on pMTL80000 universal *Clostridium* expression vectors^22^ with the addition of two GG sites flanking a *ccdB* survival gene along with *Clostridium* promoter and terminator flanking the GG sites. We also constructed pD2 and pD4 to serve as promoter-terminator donor vectors. This system of six vectors (5 donor vectors and 1 recipient vector) would allow for *in vitro* expression of genes using pD1, pD3, and pD5, followed by one-step assembly of up to six DNA parts (inserts supplied by pD1, pD2, pD3, pD4, and pD5) directly into our *Clostridium* expression vector. We note that different combinations of these vectors can be used to assemble one-gene insertions (**Supplementary Figure 1A**) or two-gene insertions (**Supplementary Figure 1B**) when fewer genes are desired. It is also possible to combine more than three genes in an expression operon by using multi-cistronic donor vectors (**Supplementary Figure 1C**).

Utilizing JGI’s gene synthesis program, the GG system was expanded to create recipient and donor vectors with varying promoters, such as P_*fdx*_^17^, P_*pta*_^*18*^, P_*pfor*_ ^*19*^, and P_*wl*_ ^*18*^. Additionally, GG sites were varied in recipient vectors to allow assembly of anywhere between two to six parts with varying promoters, resulting in a total of 58 modular vectors (**Supplementary Figure 1D**; **Supplementary Table 3**). The variety of assembly options using different DNA parts (**Supplementary Table 2**) increases the versatility of this vector system.

### Evaluation of the vector system for CFE

The highest-yielding cell-free systems take advantage of T7 RNA polymerization and are substantially affected by changes in plasmid architecture.^32,33^ For example, our previous work using the pJL1 vector that leverages T7 RNA polymerase to make mRNA achieved protein yields of ∼2.7g/L super-folder green fluorescent protein (sfGFP).^34^ In order to adapt our robust pJL1 vector for GG compatibility, we chose to test the insertion of three BsaI site designs in pJL1 (**Figure 2A**; **Supplementary Table 2**). Specifically, GG sites with sequences TCAT, AATG, or CTTA were introduced between the ribosome binding site (RBS) and start codon, which increased the spacer length between these two elements by 1 to 3 nucleotides. This created three distinct donor vectors each with three possible BsaI cut sites. We evaluated each of these nine designs in a cell-free gene expression reaction based on the PANOx-SP system to produce sfGFP to first assess the impact of GG sites on protein expression. After 20 h, cell-free reactions produced comparable or slightly greater concentrations of sfGFP than the unaltered pJL1 plasmid (**Figure 2B**). For the donor vectors to be compatible we had to choose the same variant for each of the three vectors. Thus, we chose variant 2 as the highest-performing set. Variant 2 GG vectors were validated further with expression of the enzymes phosphotransbutyrylase (Ptb) and butyrate kinase (Buk) from butyric acid metabolism in *C. acetobutylicum* ATCC 824 (**Figure 2C**). This experiment highlights the importance of genetic context for expression of different proteins, yielding variable amount of protein for Ptb and Buk despite nearly identical expression of sfGFP.

**Figure 2.**
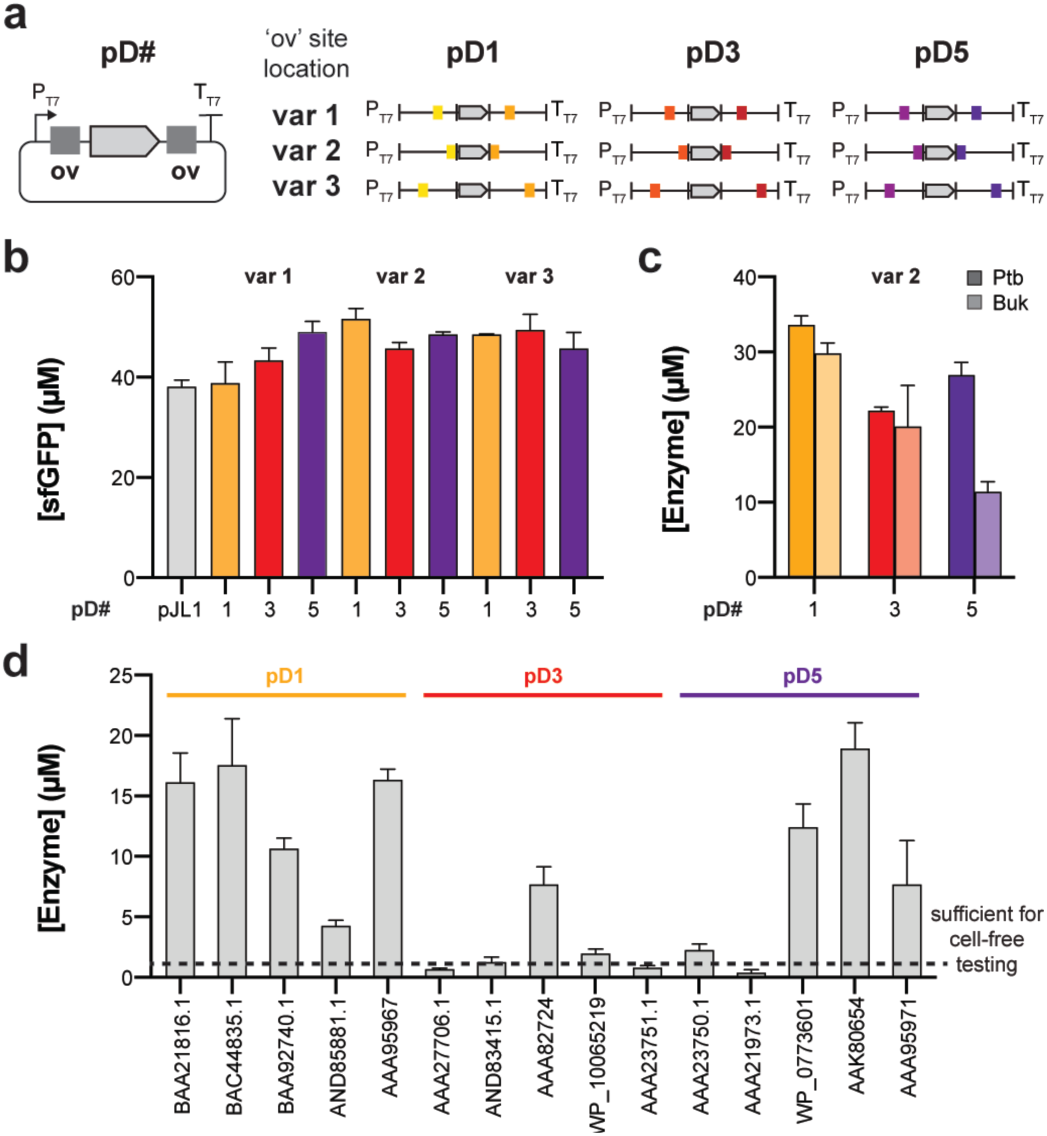
Cell-free expression of Golden Gate compatible vectors is sufficient for prototyping biosynthetic enzymes. (a) We tested three variants (change in where BsaI sites are located) of each of the three donor plasmids in CFE using a reporter sfGFP. (b) We measure sfGFP concentration by fluorescence at 20 h after reaction start. Data is shown for n=2 independent experiments with average error. (c) We switched the sfGFP reporter for Ptb and Buk in variant 2 for each of the three donor vectors and measured protein concentration at 20 h via radioactive leucine incorporation. Data is shown for n=2 independent experiments with average error. (d) 15 enzymes of interest for acid and alcohol fermentation were codon optimized for *C. autoethanogenum* and cloned into pD1, pD2, and pD3. Protein expression was measured at 20 h for n=3 independent experiments. All error bars represent 1 s.d.

We next evaluated whether *C. autoethanogenum*-optimized genes would be sufficiently expressed in our established cell-free assays. To test this, we constructed a panel of 15 *Clostridium-*optimized biosynthesis genes (**Supplementary Table 4**) related to acid/alcohol fermentation from a variety of organisms in pD1, pD3, and pD5. After a 20-h CFE reaction we observed a range of expression all significantly lower (10-fold) than what we saw for *E. coli-*codon-optimized sfGFP, Ptb, or Buk (**Figure 2D**). For enzymes that are not expressed well, separately expressing the *E. coli-*codon-optimized versions can improve enzyme expression (**Supplementary Figure 2**; **Supplementary Table 4**). Although the soluble protein yields are generally lower using *C. autoethanogenum* (31% GC content) codon-optimized sequences in *E. coli*-based (50% GC content) cell extract, these concentrations should still be sufficient for *in vitro* pathway prototyping. Expressing enzymes at concentrations greater than 1 µM provides at least 0.1 µM enzyme after dilution upon pathway assembly *in vitro* which we have found to be sufficient for prototyping.^13^ When necessary, *E. coli-*codon optimization can be used. With GG-compatible vectors at hand we next sought construct *in vivo* expression plasmids.

### Six-part DNA assembly from CFE vectors into *Clostridium* expression plasmid

Once the pD vectors were successfully validated in CFE, these modified vectors were then used for testing the efficiency of multiple-part assembly directly into a *Clostridium* expression vector with a variety of biosynthetic genes. Specifically, we carried out a six-part GG assembly that contained: (i) a recipient vector based on pMTL8315 backbone containing a promoter (P1) and terminator (T3) flanking the two GG sites (pCExpress), (ii) pD2 and pD4, both containing terminator and promoter combinations (i.e., T1-P2 and T2-P3), and (iii) pD1, pD3, and pD5 containing gene 1, gene 2, and gene 3 (**Figure 3A**; **Supplementary Table 1**). The assembly mixture was transformed into our *E. coli* cloning strain and six colonies were picked and genotyped by PCR which indicated 90-100% of the picked colonies had plasmids with all six parts correctly assembled (**Figure 3B**). These were confirmed via sequencing. The six-part assembly was validated using a different set of genes and promoter-terminator combinations for at least five additional designs (**Supplementary Table 5**). In a total of 20 manual assemblies were carried out with efficiency ranging from 70-95%.

**Figure 3.**
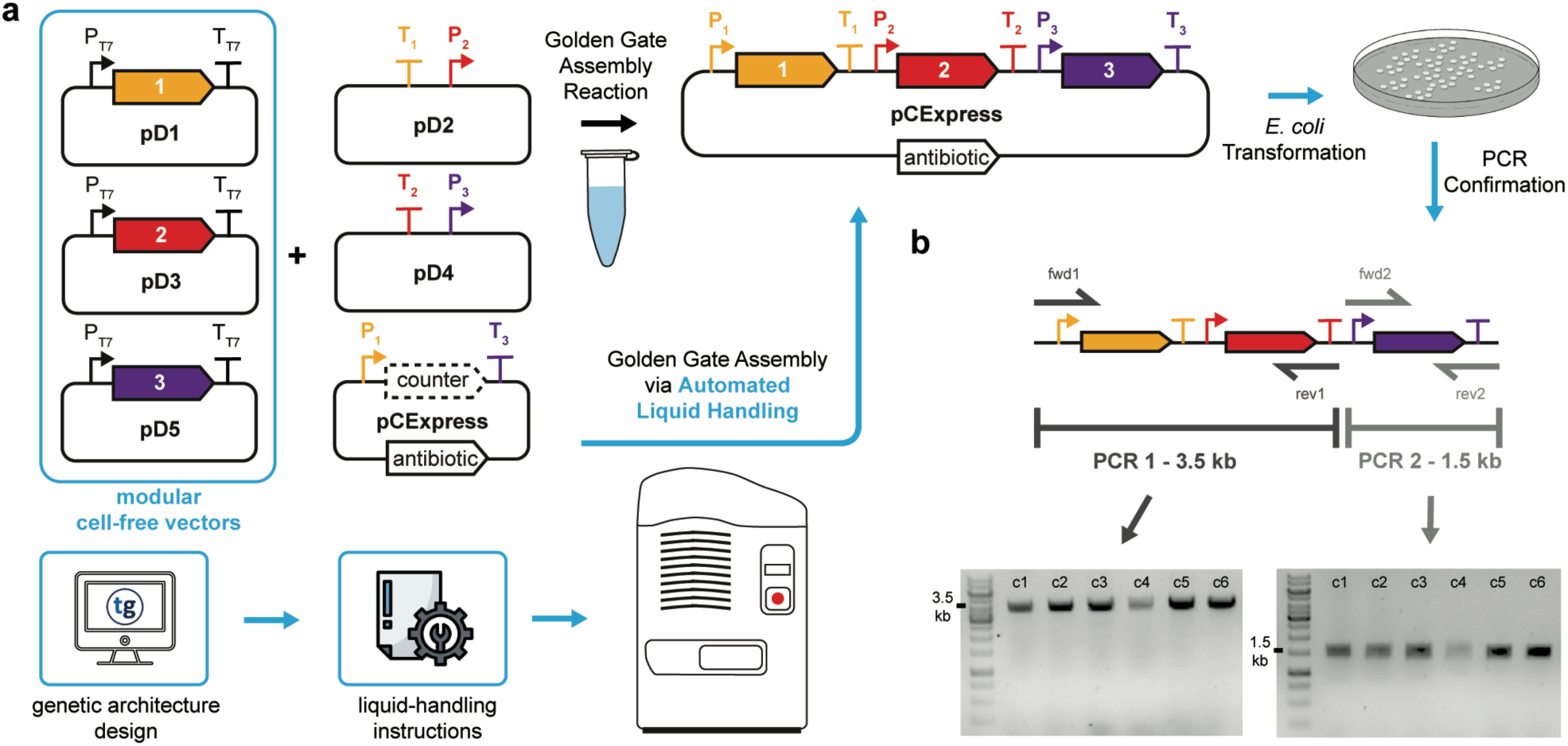
Golden Gate assembly of a 3-gene construct using compatible cell-free vector system. (a) A schematic representation of our Golden Gate assembly workflow including automated assembly consisting of computational design of plasmids, liquid-handling instructions, plasmid assembly, and plasmid confirmation. (b) PCR confirmation of plasmid assembly in six colonies containing the constructed *Clostridium* expression vector.

### Workflow automation

Workflow automation can improve throughput and reliability. CFE reactions can routinely be performed using liquid-handling robotics.^35,36^ These reactions can be scaled down to 2 µL without significant changes in protein expression.^37^ In addition, GG assembly for *in vivo* expression can also be automated. After demonstrating successful assembly of up to six DNA parts using a manual workflow, we then developed an automated workflow to increase our DNA assembly throughput (**Figure 3A**). Due to the complexity of biological systems, it is often necessary to test a large number of enzyme homologs along with different promoters to obtain an optimal engineering solution. Indeed, testing just five homologs and three promoters for a three gene operon would yield 3,375 different permutations. However, this experimental throughput is difficult and laborious when using manual techniques and procedures. Automated, well-informed designs help to increase the number of designs that can be generated, the speed these designs can be generated, and it helps to narrow down the design space prioritizing the best candidates to be built and tested, saving lab resources.^38^ In order to increase throughput, efficiency, and accuracy of our strain engineering pipeline, free up researchers from repetitive tasks, and increase results reproducibility, we validated a Golden Gate DNA assembly automated protocol on two automation systems. Both the design of constructs and the worklists to run the experiments were generated by J5 software.^39,40^ We assembled three- to six-part GG assemblies using both a Hamilton STARLet liquid-handling robot and a Labcyte Echo 525 acoustic liquid-handling robot with greater than 90% efficiencies.

## Conclusion

In this study, we describe a set of modular vectors for both cell-free gene expression and cloning into *Clostridium* expression plasmids. This framework allows facile testing of biosynthetic pathways *in vitro* and *in vivo* for shorter engineering cycles and enables an improved workflow between our *in vitro* team, our *in vivo* team, and JGI, without lengthy and costly re-synthesis and/or subcloning. The ‘Cell-free to *Clostridium*’ vector system is easy to use for Golden Gate assembly of up to six parts (three open reading frames with unique promoter and terminator sequences) at once with up to 90% efficiencies and feeds directly into the JGI’s Community Science Program platform. For longer operons, genes can be sequentially located on each of the CFE vectors (pD1, pD3, and pD5). These vectors along with laboratory automation have already increased the speed and efficiency of our workflows and will continue to facilitate the ability to prototype biosynthetic pathways *in vitro* followed by *in vivo* cloning pipelines. Standardization of these vector systems allows for new simplified workflows. The pJL1 cell-free vector and variants thereof are routinely used in multiple bacterial cell-free systems (i.e., *E. coli*,^19^ *Clostridium*,^10^ *Pseudomonas*,^23^ *Streptomyces*,^24,41^ *Vibrio natriegens*^42,43^). In addition, the pMTL vector system has been demonstrated in several Clostridia species (i.e., *autoethanogenum, ljungdahlii, acetobutylicum, beijerinckii, difficile, sporogeneses, perfringens, pasteurianum, tyrobutyricum*) as well as other Gram-negative and Gram-positive model organisms such as *E. coli* and *Bacillus*.^22,44^ Taken together, the breadth of bacterial cell-free systems that can use the pJL1 vector and ubiquity of Golden Gate cloning suggests broad applicability of our plasmid vector system. Looking forward, we anticipate this system of vectors will allow researchers to integrate more *in vitro* prototyping practices into their existing workflows across multiple organisms to speed up metabolic engineering efforts.

## Acknowledgements

We would like to acknowledge Jan-Fang Chen, Robert Evans, Yasuo Yoshikuni, and Miranda Harmon-Smith from the U.S. Department of Energy Joint Genome Institute, a DOE Office of Science User Facility, for their DNA synthesis, DNA assembly, and support in this work as part of the Community Science Program (CSP).

## Funding

This work was supported by the U.S. Department of Energy (DOE) Biological and Environmental Research Division (BER), Genomic Science Program (GSP) for funding of this project under Contract No. DE-SC0018249. We also thank the Joint Genome Institute Community Science Program Project 503280. The work conducted by the U.S. Department of Energy Joint Genome Institute, a DOE Office of Science User Facility, is supported by the Office of Science of the U.S. Department of Energy under Contract No. DE-AC02-05CH11231. We also thank the following investors in LanzaTech’s technology: BASF, CICC Growth Capital Fund I, CITIC Capital, Indian Oil Company, K1W1, Khosla Ventures, the Malaysian Life Sciences, Capital Fund, L. P., Mitsui, the New Zealand Superannuation Fund, Novo Holdings A/S, Petronas Technology Ventures, Primetals, Qiming Venture Partners, Softbank China, and Suncor.

## Conflicts of Interest

A provisional patent has been filed on this vector system (US 62/943,036).

## Author Contributions

A.S.K., F.E.L., A.J., M.K., and M.C.J. conceived the study. A.S.K., B.V., and B.J.R. performed cell-free experiments. F.E.L., S.G., A.G., and M.P. performed *in vivo* experiments. S.D.S., M.K., and M.C.J. provided supervision. All authors wrote the manuscript.

## Supplementary Information for

**Supplementary Table 1.**
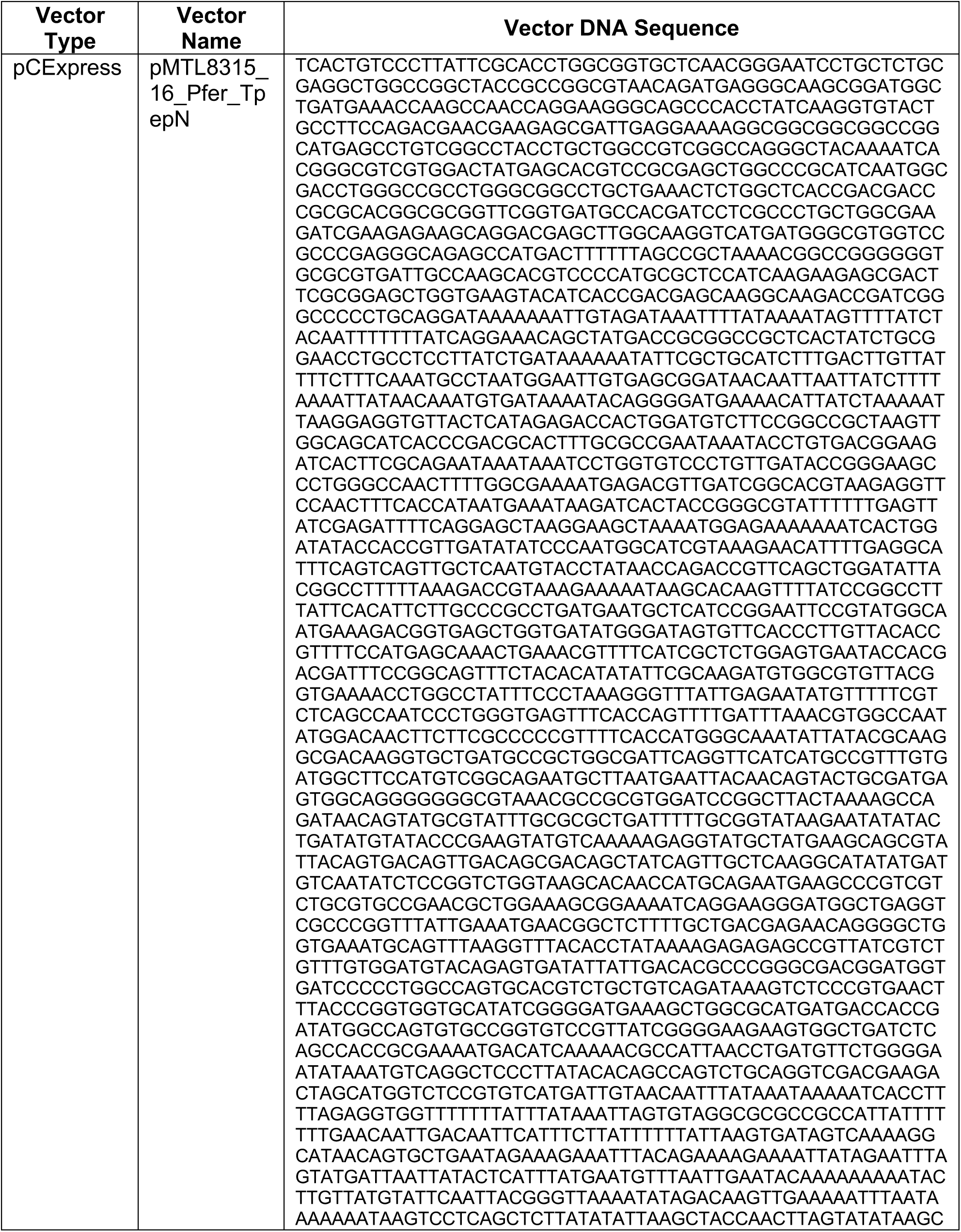
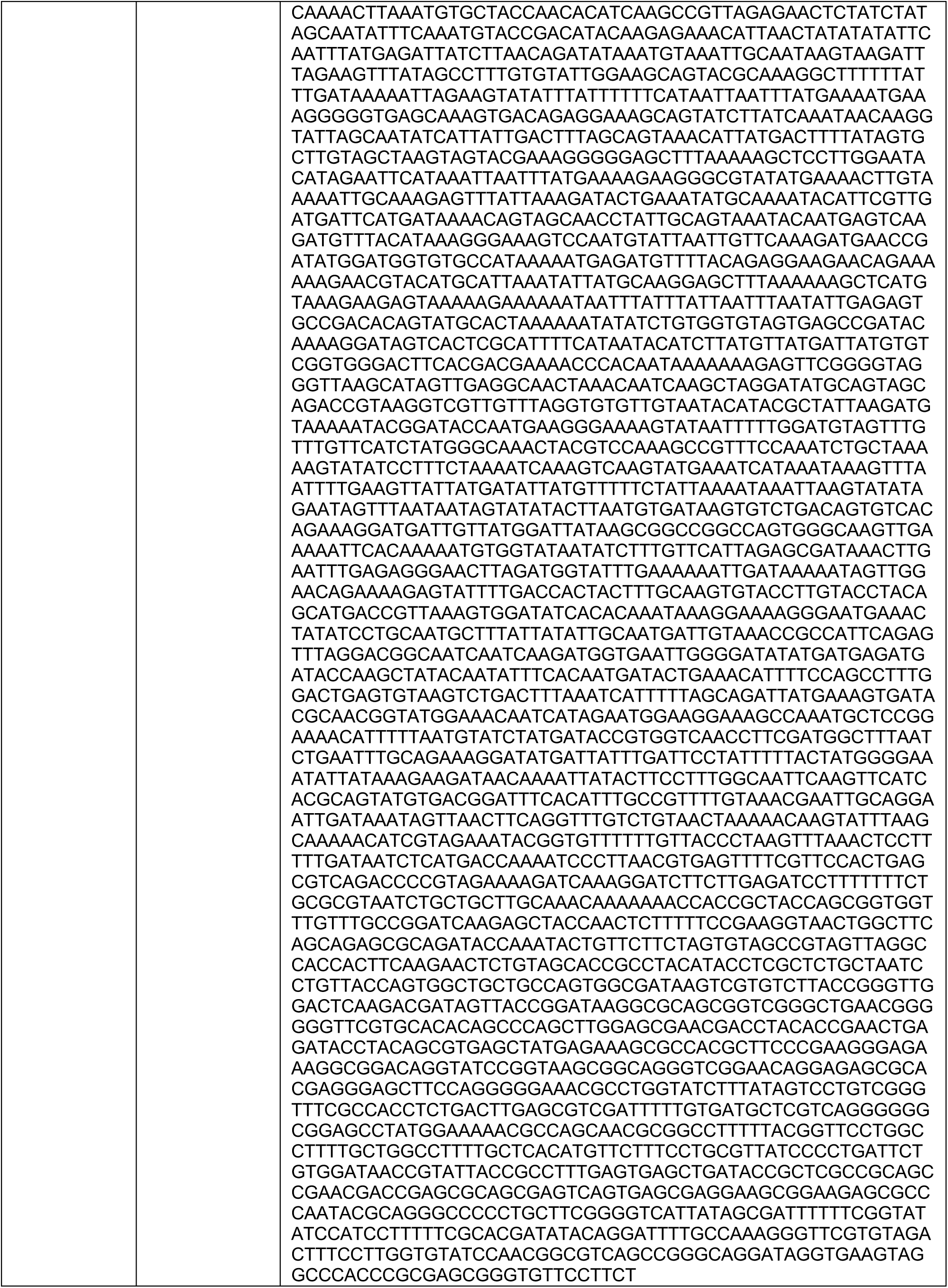
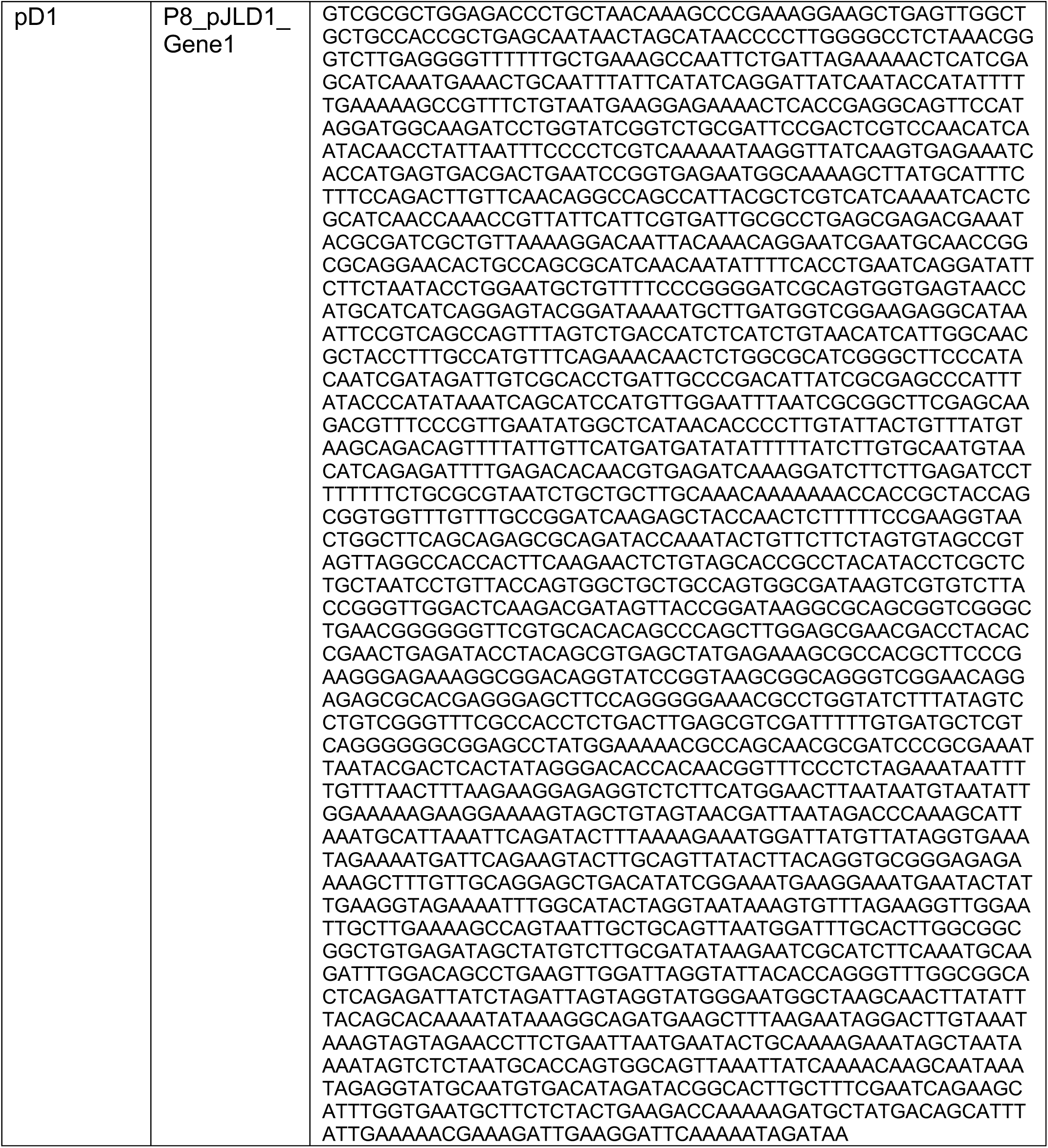
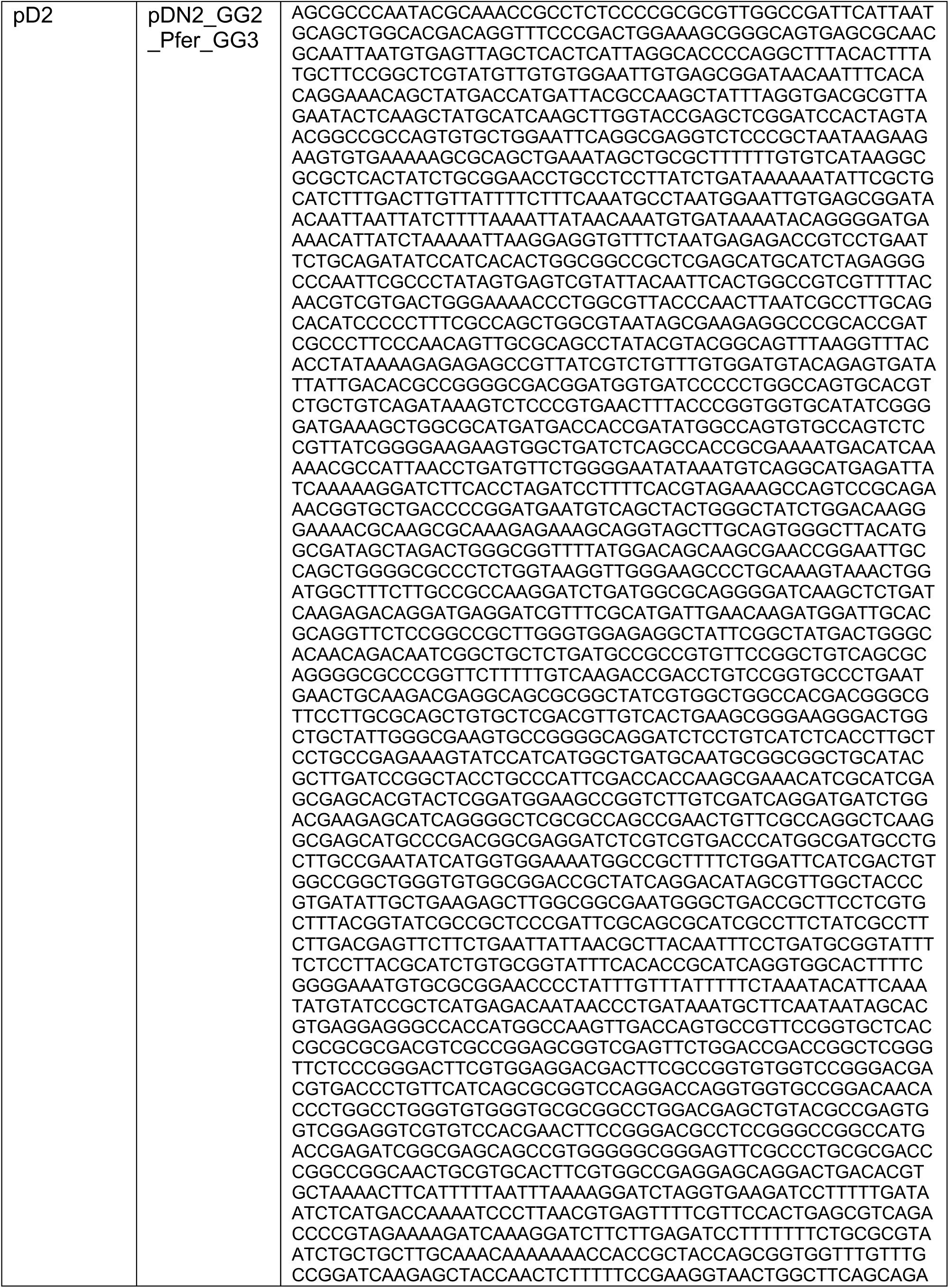
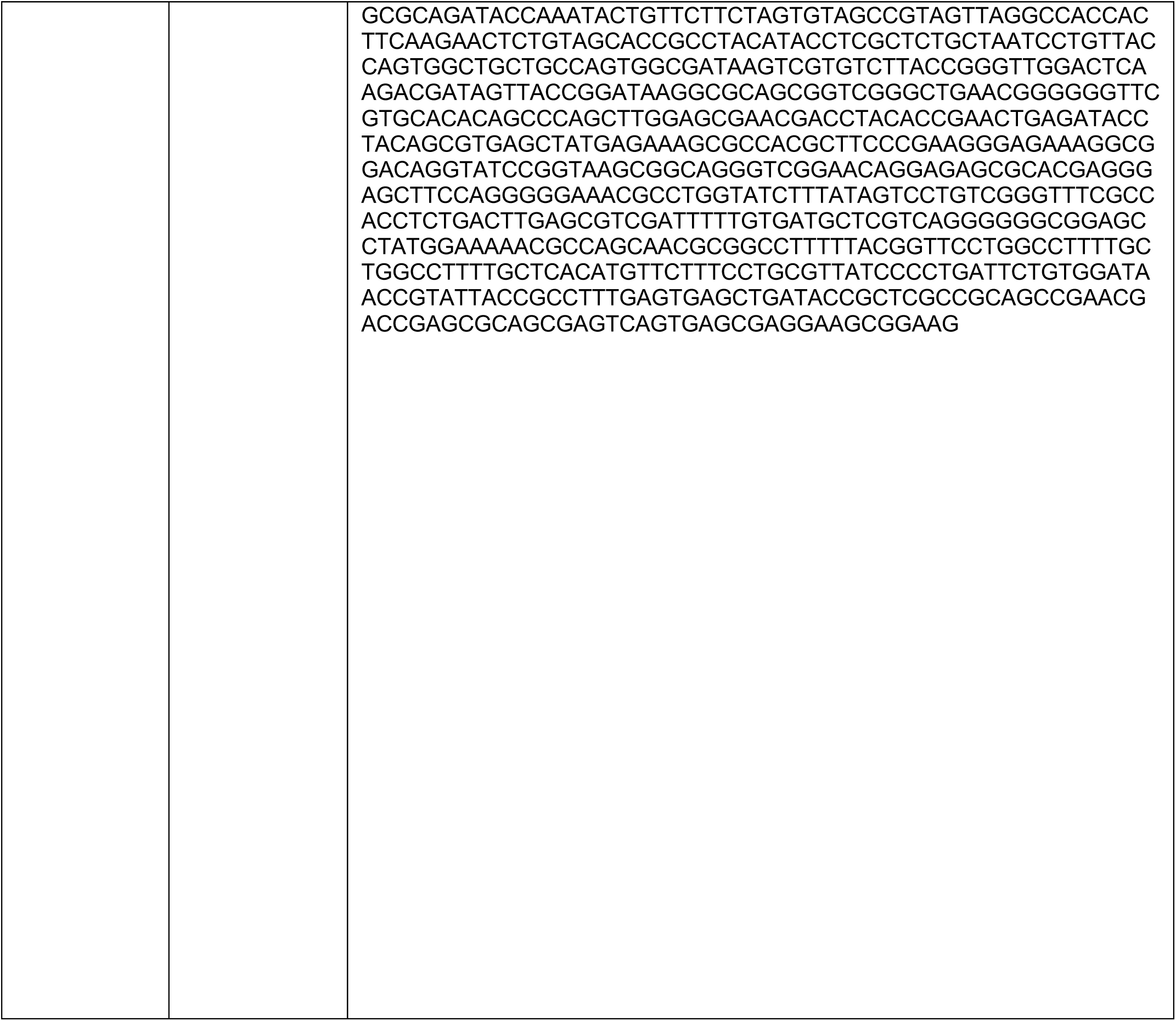
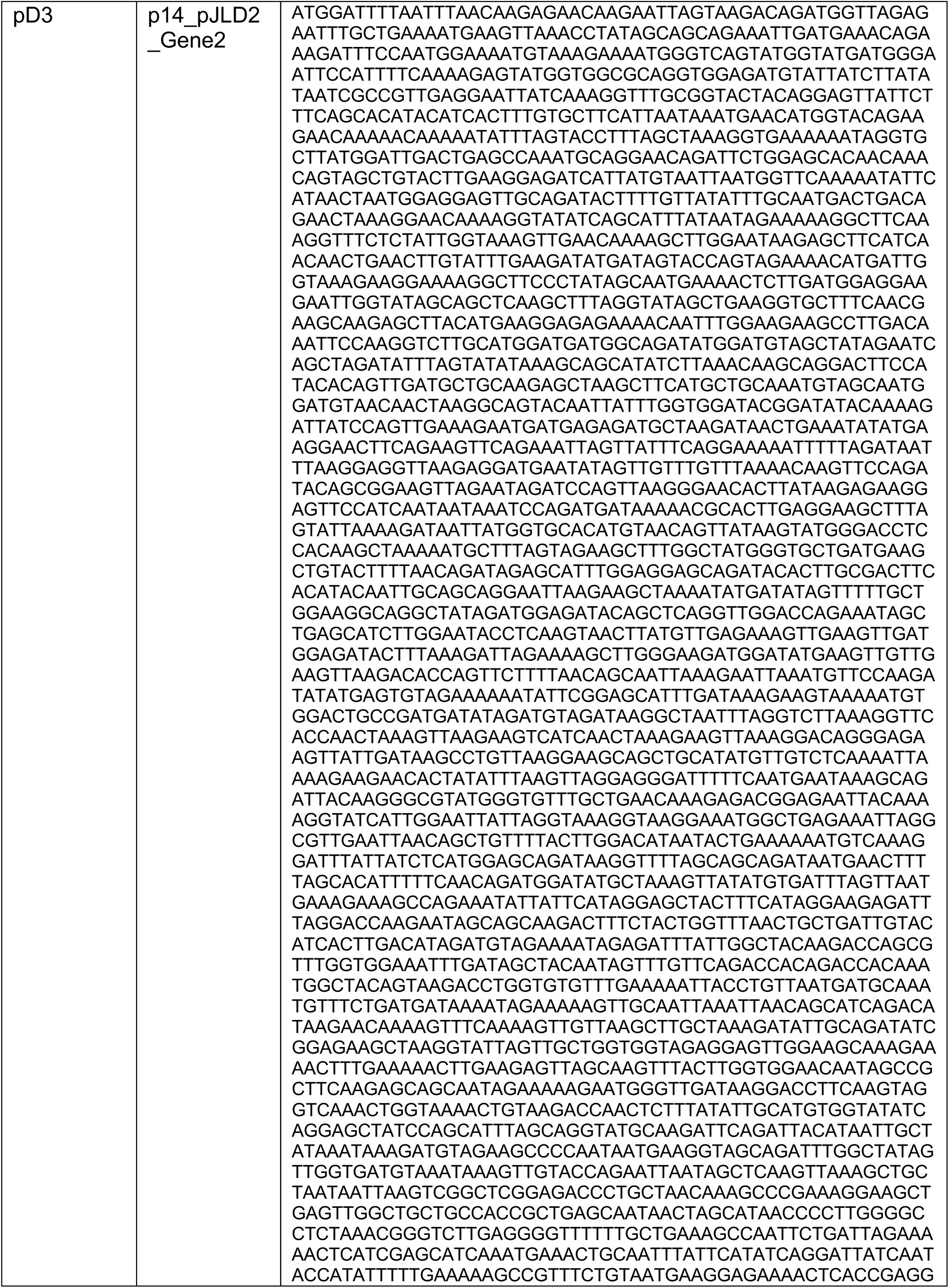
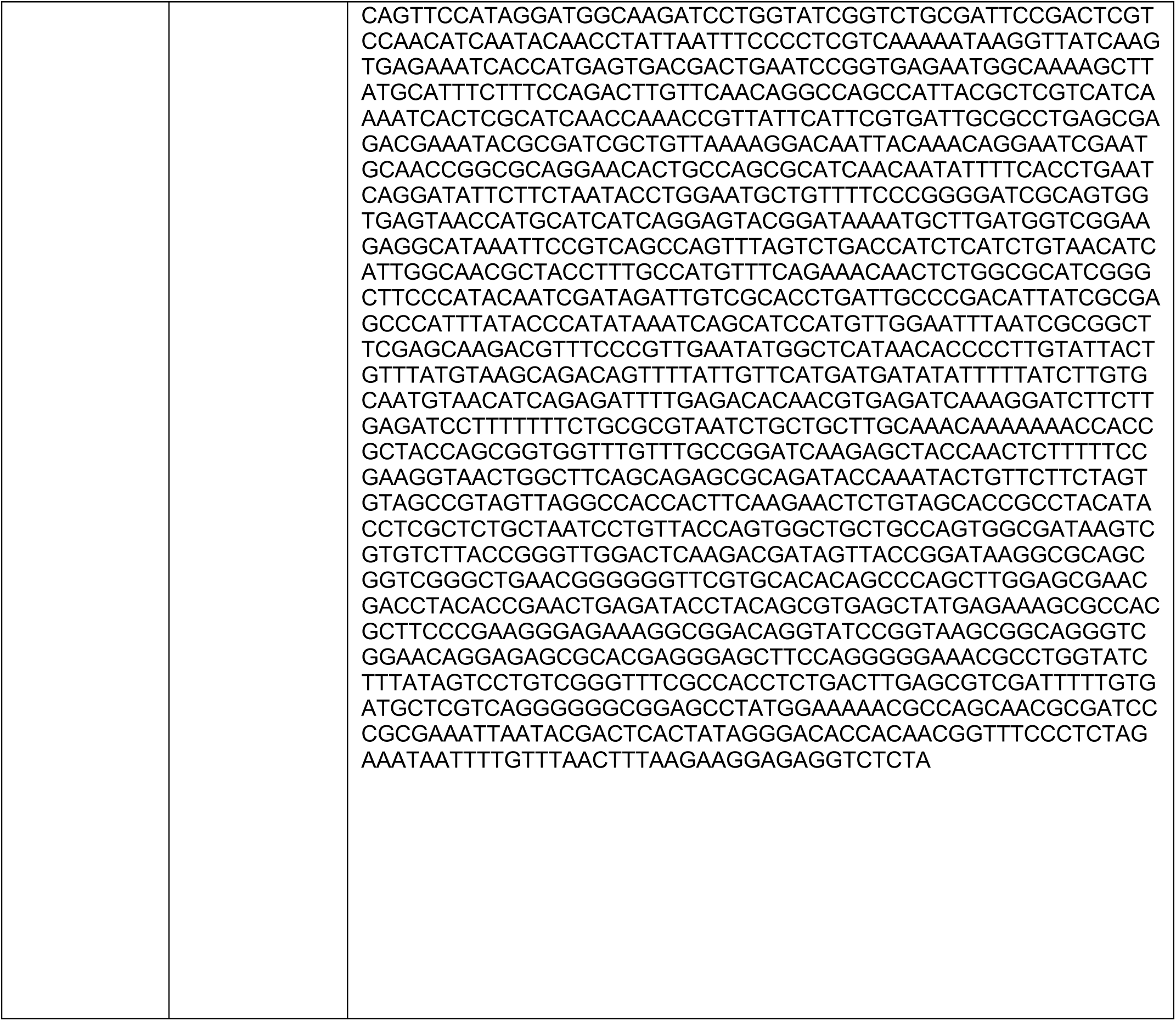
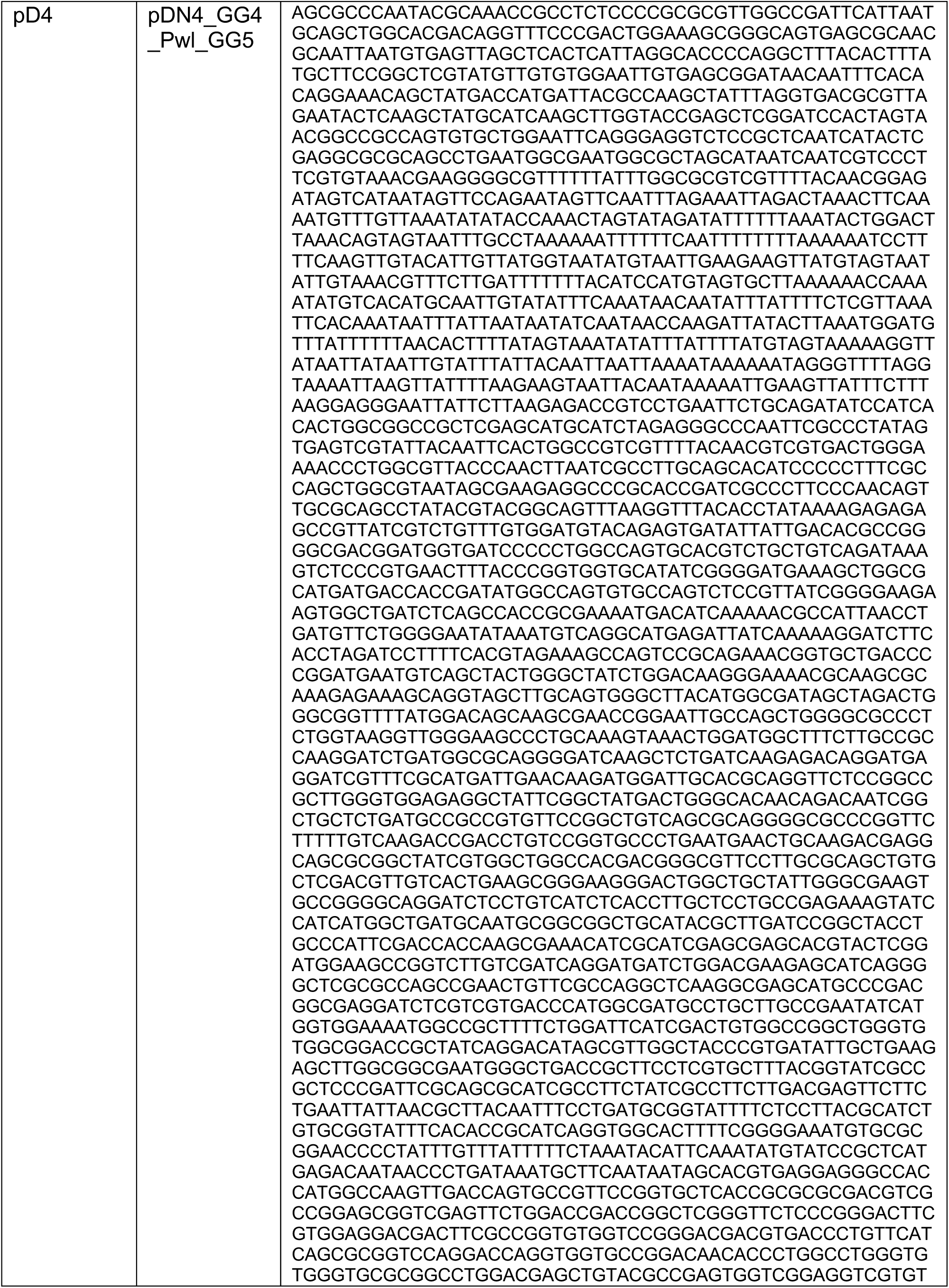
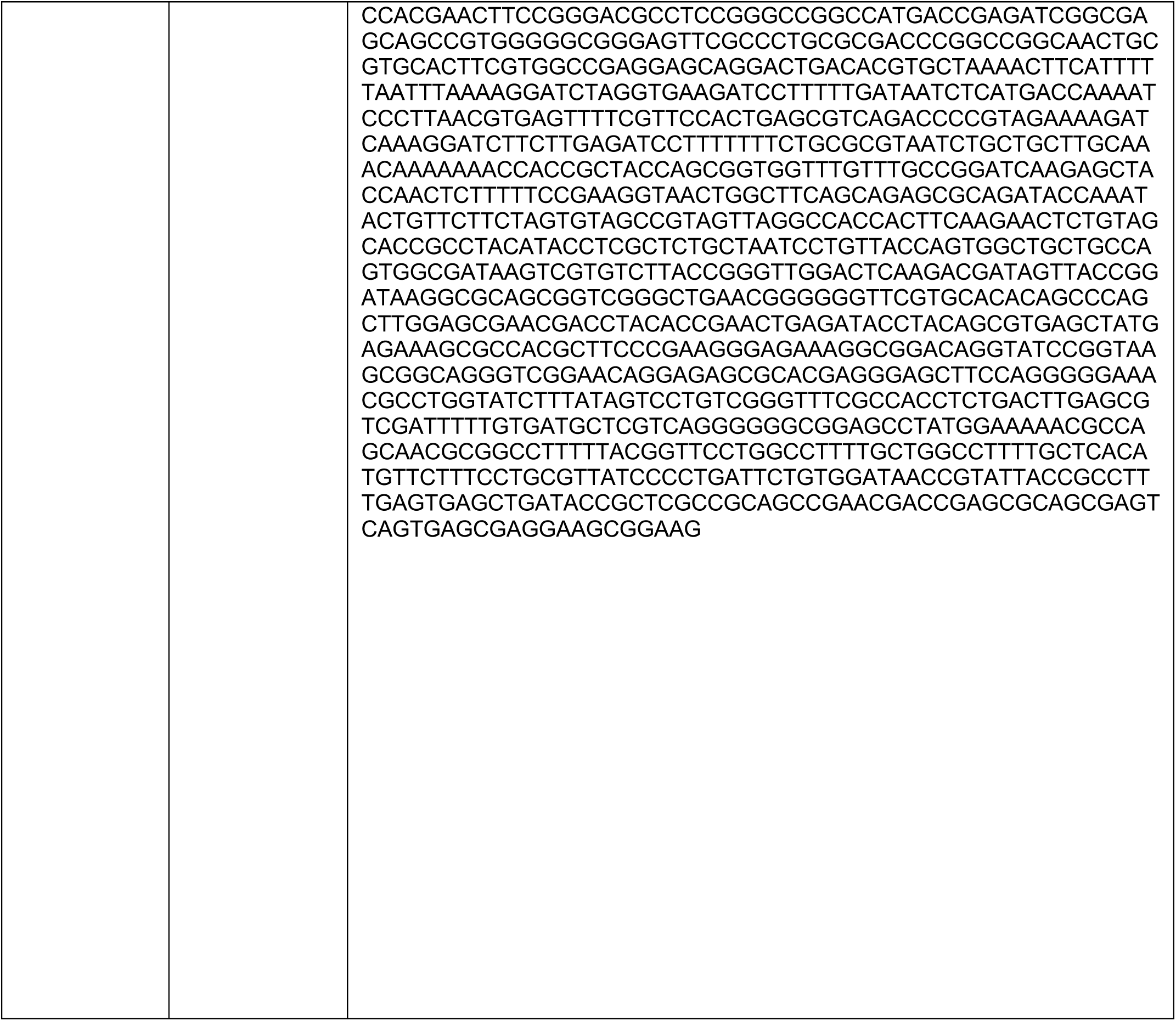
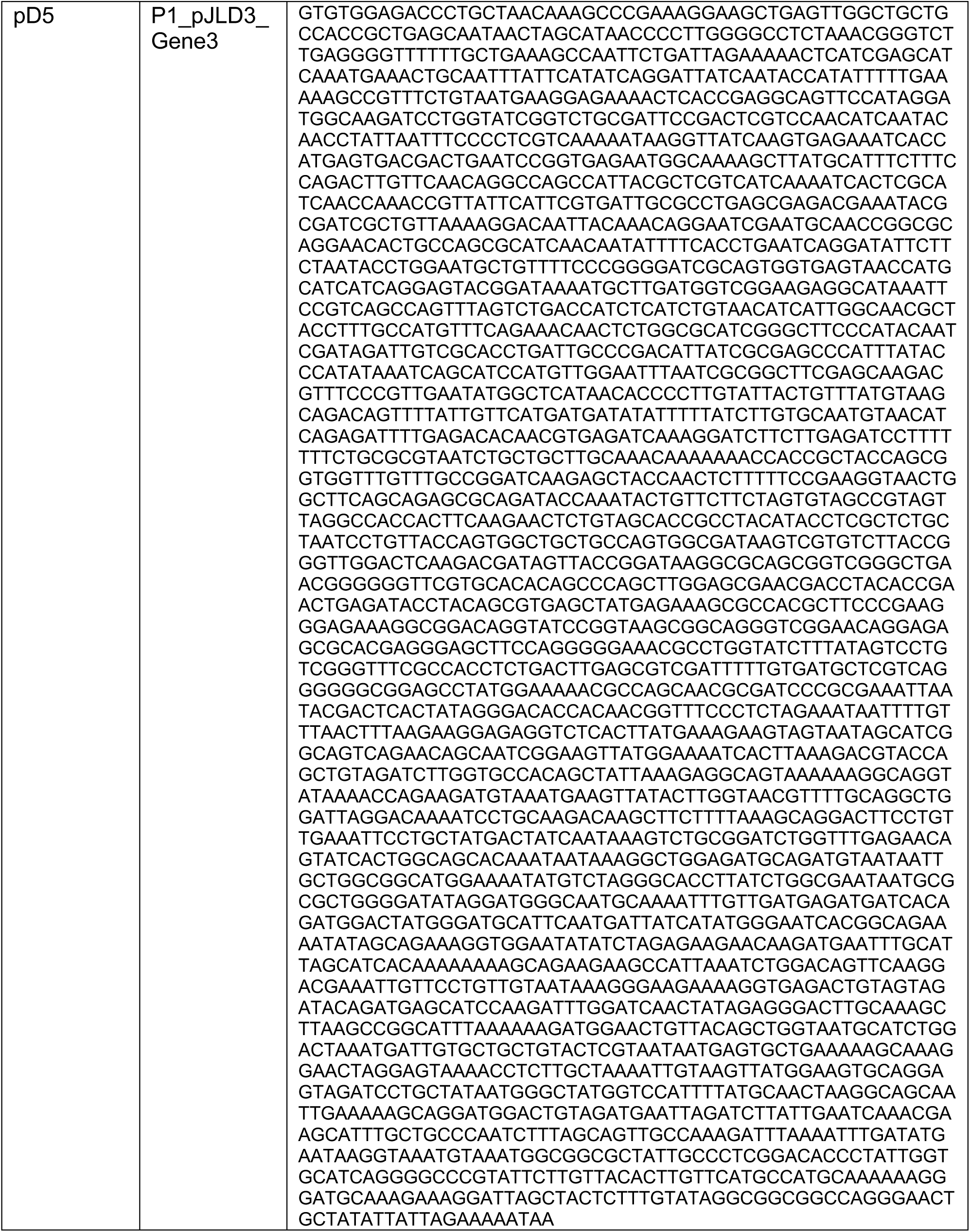
DNA Vector Sequences. Below is a table of all vectors used in this study. The vector type is what is referenced throughout the manuscript.

**Supplementary Table 2.**
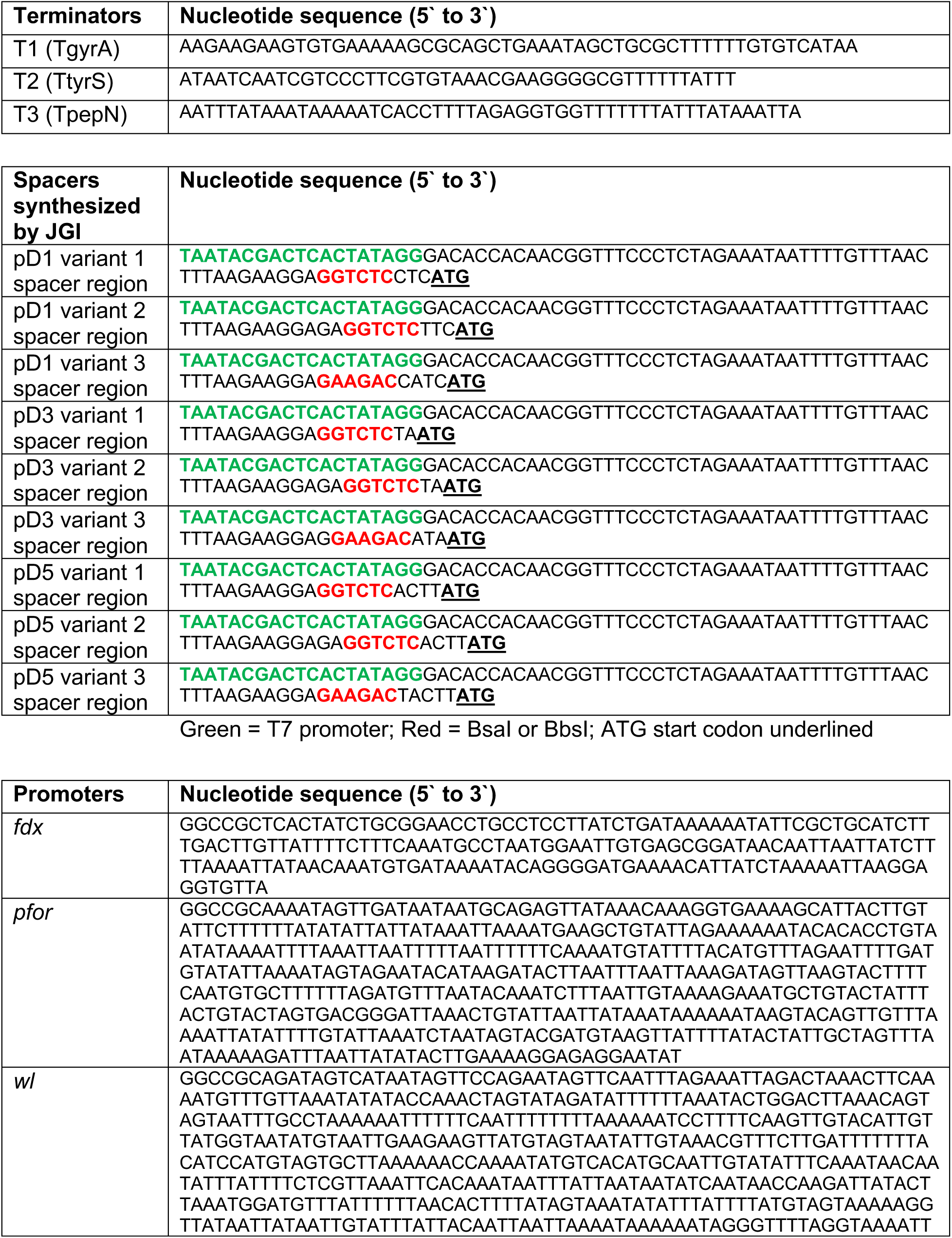
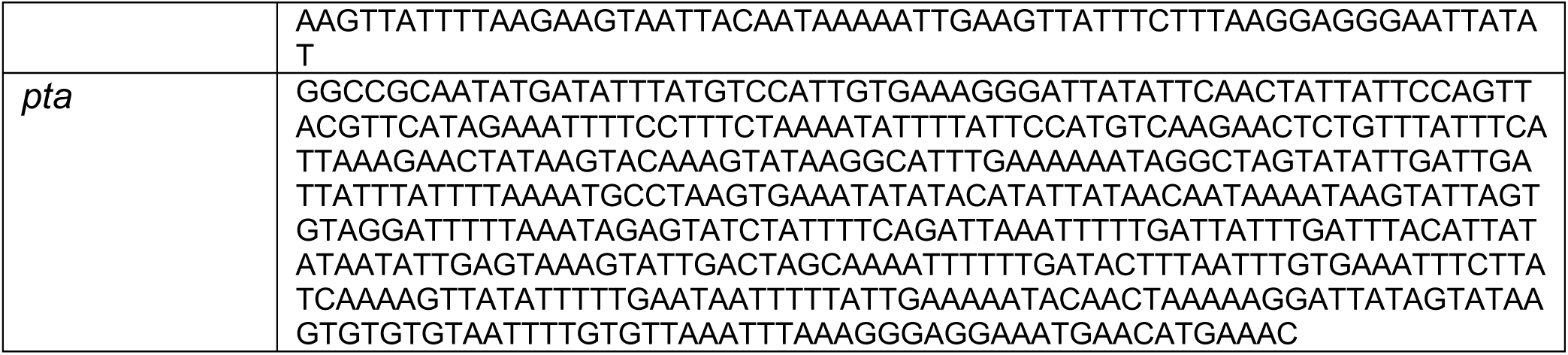
List of DNA Parts. Terminators, spacers, and promoters are used to construct operons in pCExpress vectors used in this study are listed.

**Supplementary Table 3.**
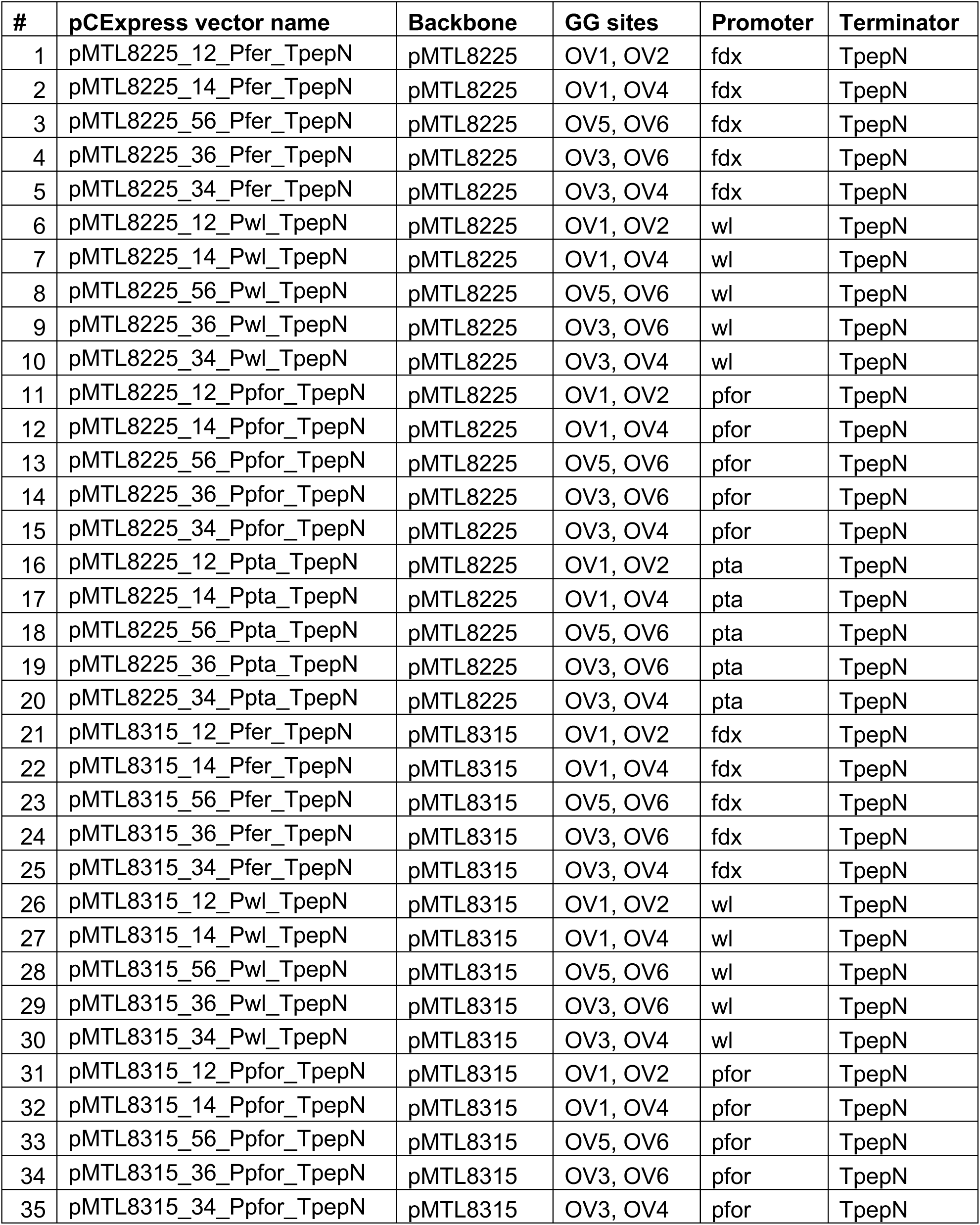
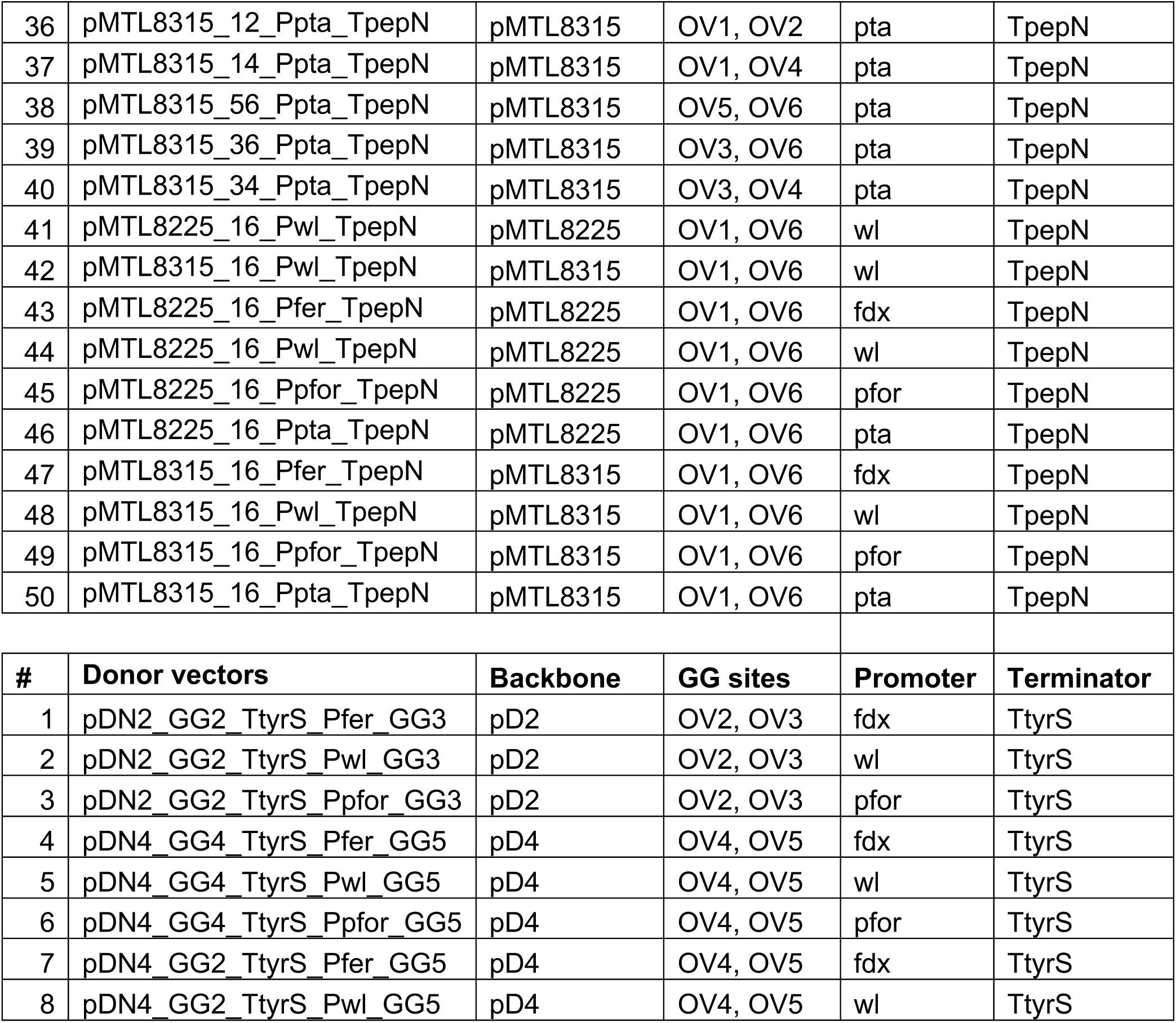
List of pCExpress and pD2/pD4 variants for assembly versatility. Each variant is listed with name and backbone vector is derived from along with which overhangs, promoter, and terminator is present.

**Supplementary Table 4.**
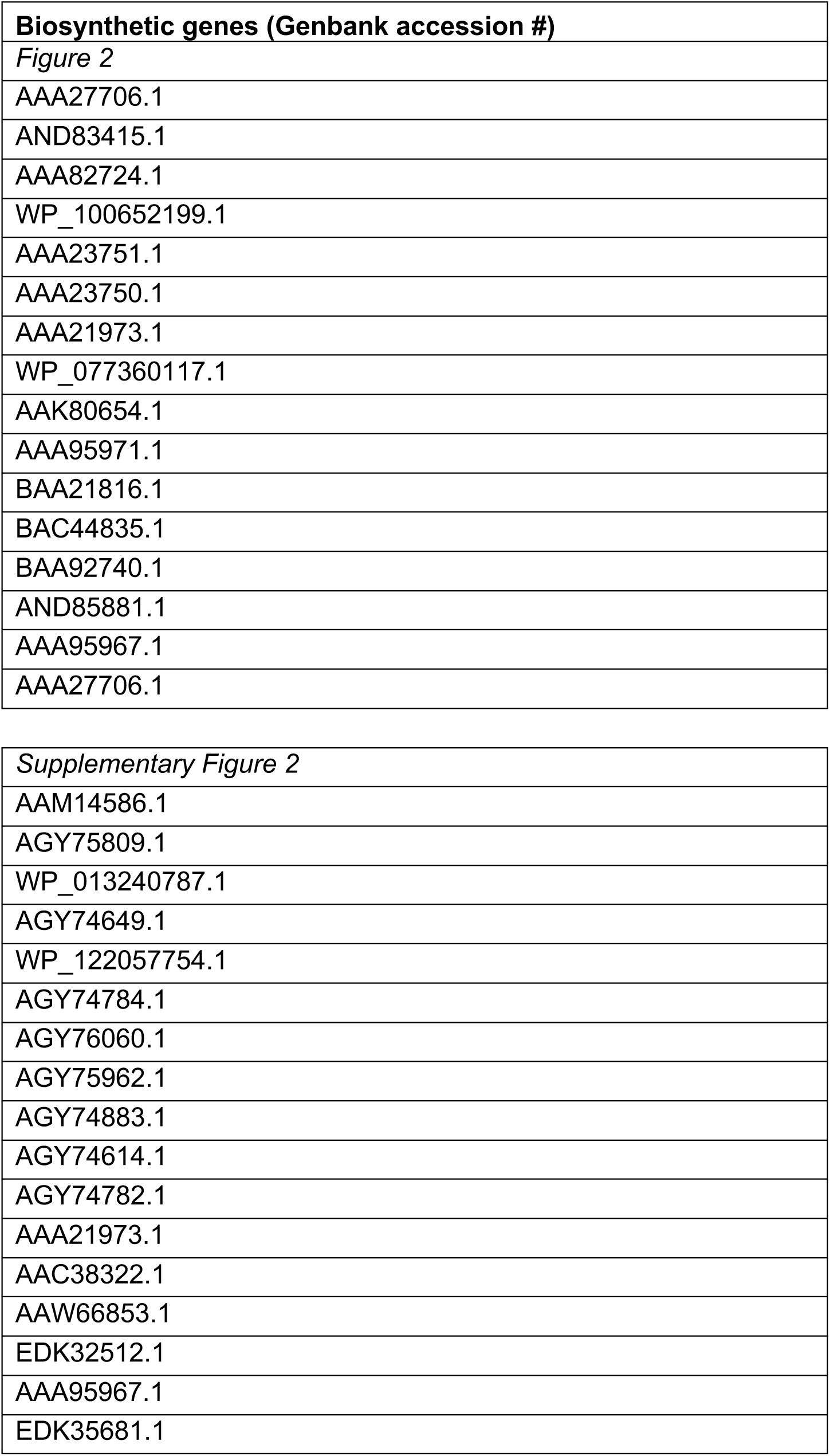
List of biosynthetic genes used CFE experiments. Genes used are represented by their Genbank accession numbers.

**Supplementary Table 5.**
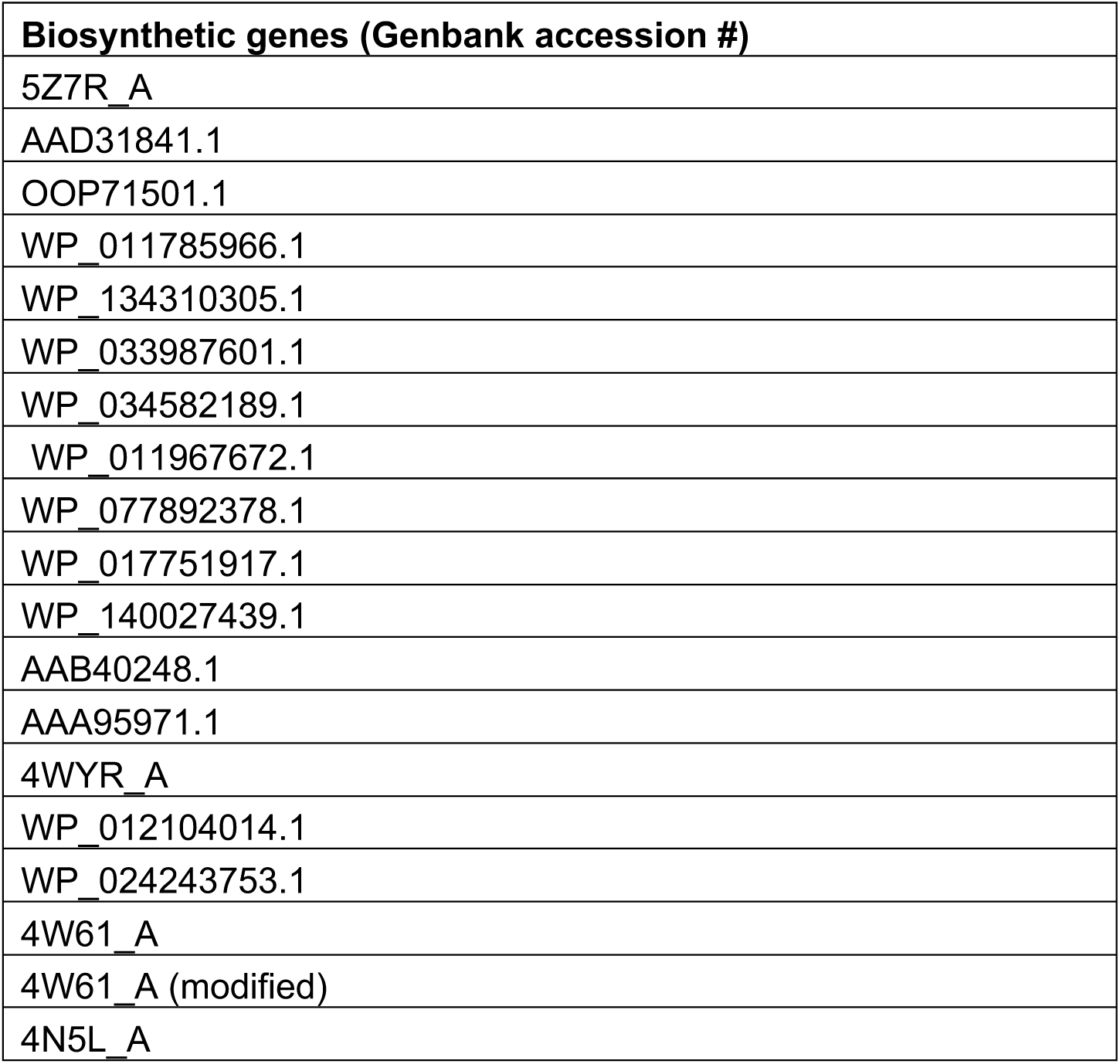
List of biosynthetic genes used to assess codon usage and assembly efficiency. Genes used are represented by their Genbank accession numbers.

**Supplementary Figure 1.**
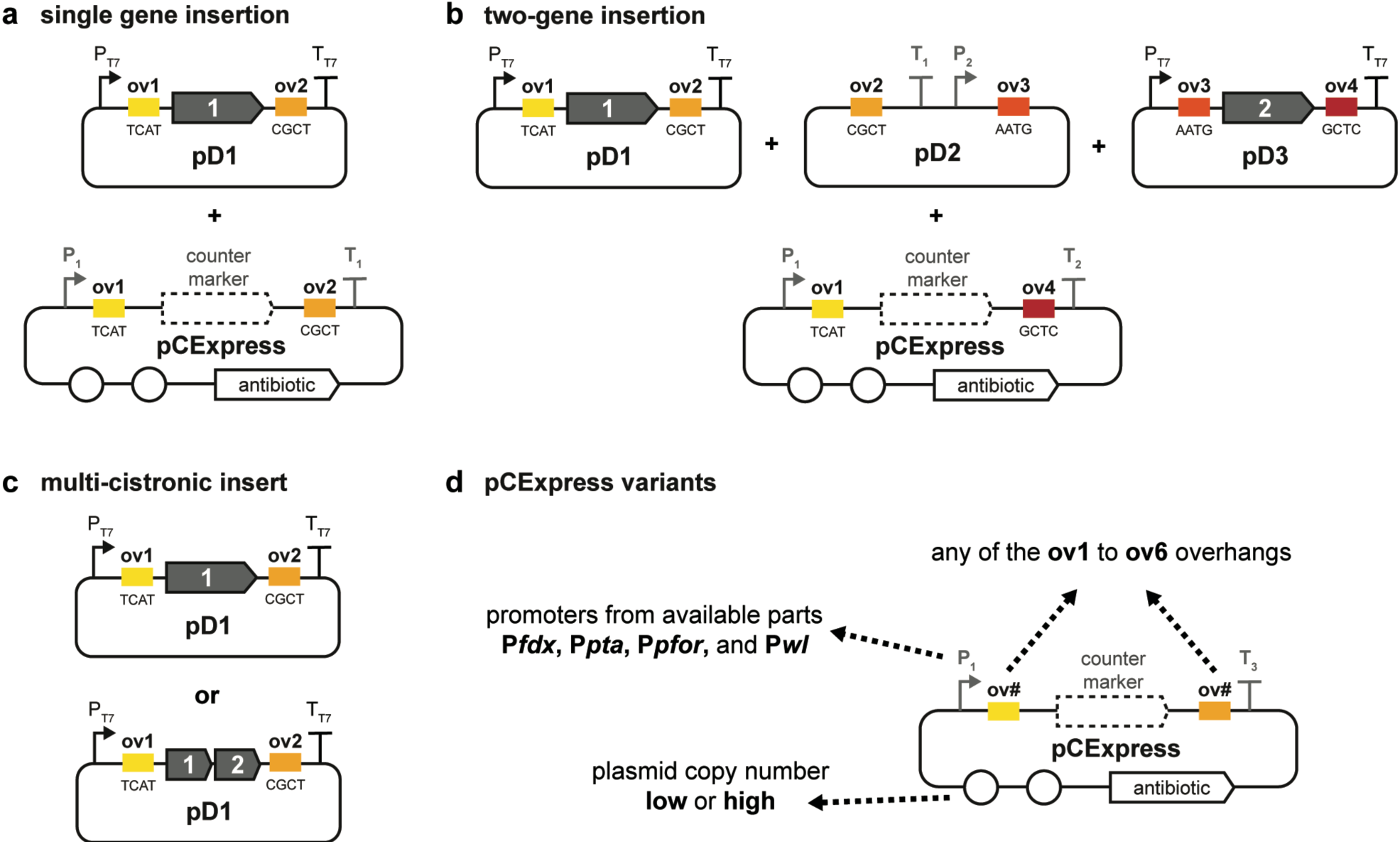
A modular ‘cell-free to *Clostridium*’ vector system. (A) Two-part assembly for a single gene insertion. (B) Configuration of donor vectors that enables two-gene insertion using defined Ov sites. (C) Modifying pD1 (or pD2, pD3) to hold more than 1 gene allows for complete assembly of more than three genes using the ‘Cell-free to *Clostridium*’ vector system. (D) pCExpress can be varied for different assembly types. The key parameters are mentioned here with the full table of variants in **Supplementary Table 3**.

**Supplementary Figure 2.**
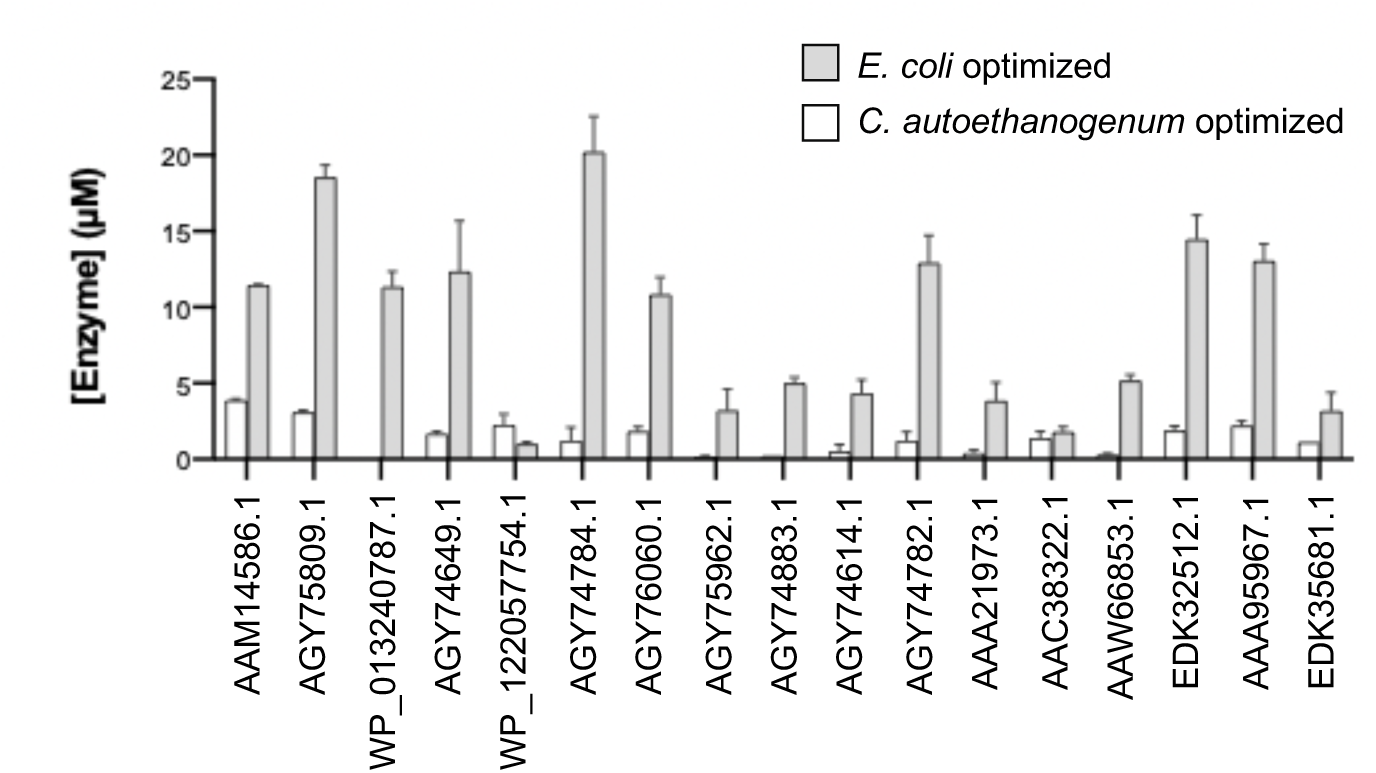
Cell-free expression of *E. coli* optimized DNA sequences produce more protein than *Clostridium* optimized sequences. 17 gene sequences were codon optimized for *E. coli* and *C. autoethanogenum* and placed in pJL1. Each was expressed in CFE with n=2. Average expression is shown after 20 h with error bars representing average error.

